# Mitotic gene conversion tracts associated with repair of a defined double-strand break in *Saccharomyces cerevisiae*

**DOI:** 10.1101/132167

**Authors:** Yee Fang Hum, Sue Jinks-Robertson

## Abstract

Mitotic recombination between homologous chromosomes can lead to loss-of-heterozygosity (LOH), which is an important contributor to human disease. In the current study, a defined double-strand break (DSB) on chromosome IV was used to initiate LOH in a yeast strain with sequence-diverged chromosomes. Associated gene conversion tracts, which reflect the repair of mismatches formed when diverged chromosomes exchange single strands, were mapped using microarrays. LOH events reflected two broken chromosomes, one of which was repaired as a crossover and the other as a noncrossover. Gene conversion tracts associated with individual crossover and noncrossover events were similar in size and position, with half of the tracts unexpectedly mapping to only a single side of the initiating break. Although the molecular features of DSB-initiated events generally agree with those predicted by current models of homologous recombination, there were unexpected complexities in associated gene conversion tracts.

## Introduction

Genomic DNA is subject to attack by a variety of endogenous and exogenous DNA damaging agents, among which are reactive oxygen species, methylating agents, crosslinking agents (e.g. aldehydes and UV light) and ionizing radiation (Hoeijmakers, 2009). When only one strand of duplex DNA is damaged, lesion removal by an excision-repair pathway creates a gap that can be filled using the complementary strand as a template. The repair of DNA damage that affects both strands is more problematic, however, and double-strand breaks (DSBs) are a particularly toxic lesion. DSBs can be repaired by homologous recombination (HR), an error-free process that uses either a sister chromatid or homologous chromosome as a template to restore the broken molecule. Alternatively, broken ends can be directly ligated by the non-homologous end joining pathway, which is generally considered an error-prone process due to the deletion/addition of sequence at the repair junction (Symington & Gautier, 2011). Defects in DSB repair lead to genome instability and are associated with cancer predisposition syndromes in humans (Tubbs & Nussenzweig, 2017).

HR that uses a sister chromatid as a repair template is typically of no genetic consequence, while that involving a homologous chromosome can result in loss of heterozygosity (LOH). LOH can reflect the non-reciprocal transfer of information from the repair template to the broken molecule, which is referred to as gene conversion (GC). Alternatively, LOH can result from reciprocal crossing over between homologous chromosomes. Crossover (CO)-associated LOH was first proposed by Stern in 1936 to explain the phenomenon of “twin spotting” in a phenotypically wild-type Drosophila strain that was heterozygous at linked loci (Stern, 1936). Specifically, when the heterozygous *yellow* and *singed* markers were in repulsion on the same chromosome arm, a rare patch of tissue with singed, short bristles was seen adjacent to a patch of yellow tissue. It was proposed that a reciprocal CO event had occurred between homologs after chromosome replication (i.e., involving only two of the four chromatids) and that the CO was located centromere proximal to both loci. Segregation of the CO chromatids into different daughter cells would then yield one daughter homozygous for the mutant *singed* marker, and the other homozygous for the mutant *yellow* marker. It should be noted that crossing over between homologs prior to replication results in four CO chromatids and subsequent segregation maintains heterozygosity, even though marker linkage is altered. In humans, mitotic LOH leads to the uncovering of recessive markers associated with human disease and with cancer.

Current molecular models of HR are largely based on meiotic and mitotic studies done in *Saccharomyces cerevisiae* (reviewed in Symington, Rothstein, & Lisby, 2014). During meiosis, physiological DSBs are produced throughout the genome by the Spo11 protein (Keeney, Giroux, & Kleckner, 1997);; during mitosis, pathological DSBs arise through replication fork collapse or as a result of spontaneous DNA damage. The production of a single, defined DSB using the HO or I-*Sce*I meganuclease has been a particularly powerful tool for examining the fate of mitotic DSBs (Haber, 1995). Following enzyme-mediated DSB induction, the 5′ ends are resected to yield 3′ tails that invade a homologous donor sequence (Figure 1). Pairing between complementary strands from different duplexes creates a region of heteroduplex DNA (hetDNA;; yellow boxes in Figure 1) and displaces the strand with the same sequence as the invading strand, resulting in a displacement (D)-loop intermediate. Extension of the invading end expands the D-loop, which exposes sequences complementary to the 3′ tail on the other side of the break. In the classic DSB repair model, the second end is captured by the D-loop, creating an additional region of hetDNA and leading to the formation of a double Holliday junction (HJ). These junctions can be cleaved to generate a crossover or a noncrossover (NCO) outcome, or can be dissolved to produce only NCOs. As an alternative to second-end capture, the D-loop intermediate can be dismantled and the extended end annealed to the 3′ tail on the other side of the DSB (the synthesis-dependent strand-annealing or SDSA pathway). This removes the D-loop associated hetDNA tract formed on the invading side of the DSB, creates a new hetDNA tract on the opposite (annealing) side of the break, and generates only NCO products. The underlying molecular mechanisms of NCO formation can be inferred by examining hetDNA-tract positions in a mismatch-repair (MMR) defective background.

**Figure 1.**
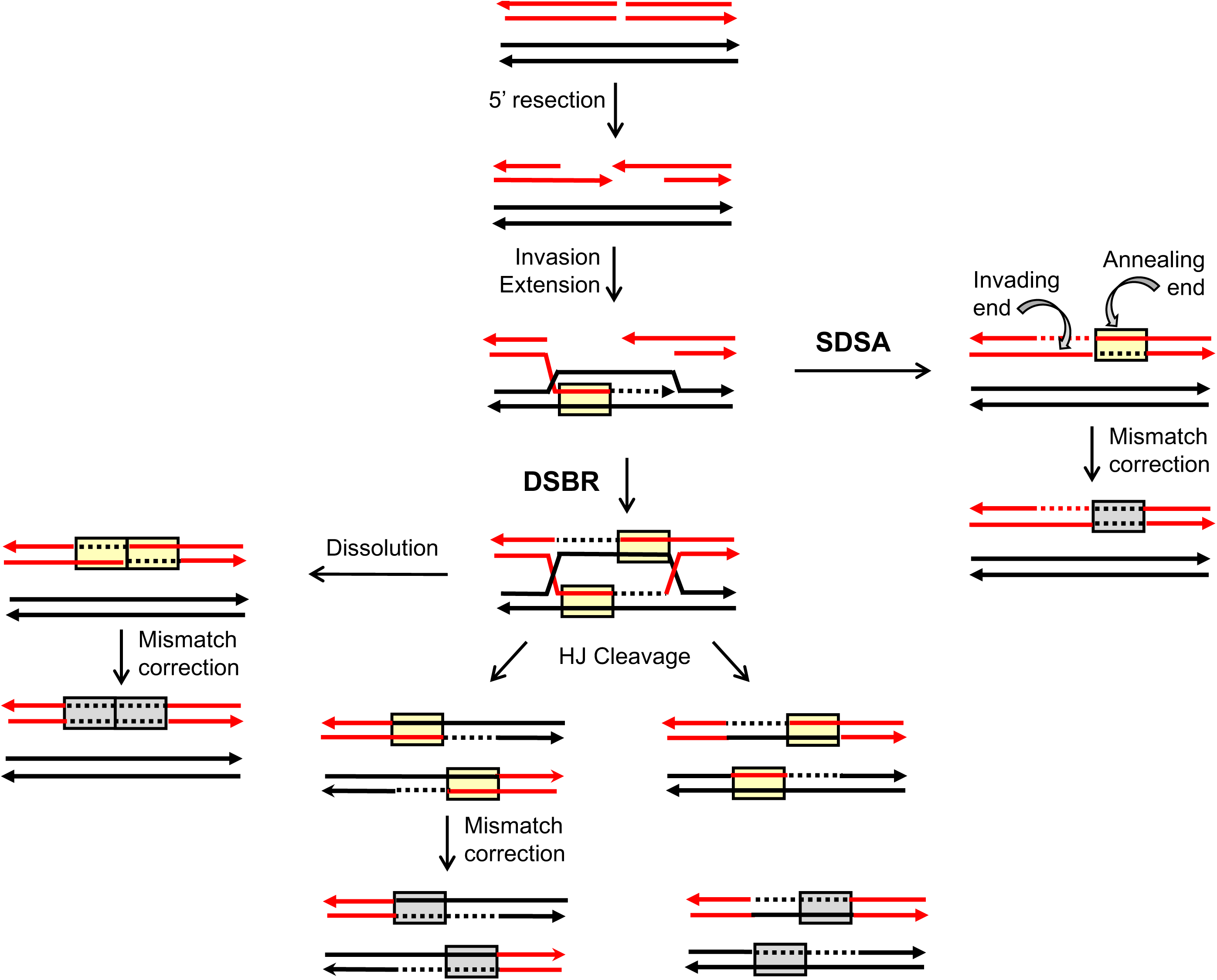
Homologous recombination models. Red and black lines represent single DNA strands and 3′ ends are marked with arrowheads; hetDNA and the GC tracts created by hetDNA correction are in yellow and gray boxes, respectively. The 5′ ends of the DSB are resected to generate 3′ single-strand tails, one of which invades the black repair template and creates a hetDNA tract. The resulting D-loop is expanded by extension of the 3′ end and is either dismantled (SDSA pathway) or captured by the 3′ end on the other side of the DSB (DSBR pathway). SDSA removes the original hetDNA tract and a new tract is created when the displaced 3′ tail anneals to the other side of the DSB. Repair of the hetDNA yields a GC tract on the annealing side of the DSB. DSBR is associated with a hetDNA tract on each side of the DSB. Cleavage of the HJs results in either CO or NCO products, and each product contains a hetDNA tract. dHJ dissolution generates a NCO, and both hetDNA tracts are on the molecule that suffered the DSB. hetDNA correction following HJ cleavage or dissolution yields a continuous GC tract that spans the site of the DSB.

Mismatches contained within HR-associated hetDNA are efficiently repaired by the MMR machinery. In meiosis, repair is biased near the initiating DSB so that the strand contributed by the broken (recipient) molecule is excised and replaced with donor information, resulting in a GC event (gray boxes in Figure 1; (Detloff, White, & Petes, 1992). Repair that occurs in the other direction removes mismatches contributed by the donor duplex, resulting in genetically silent restoration of recipient sequence. Although mitotic GC tracts are used as a proxy for the hetDNA formed during HR, their precise relationship to hetDNA has not been determined. The molecular mechanisms of NCO formation are thus most accurately inferred by mapping hetDNA tracts in an MMR-defective background. Our previous analyses of HR-associated hetDNA in a plasmid-chromosome system and in an ectopic chromosomal assay indicate that most NCOs are the result of SDSA, and that HJ cleavage generates only CO products (Guo, Hum, Lehner, & Jinks-Robertson, 2017; Mitchel, Zhang, Welz-Voegele, & Jinks-Robertson, 2010). In addition, hetDNA tracts associated with COs are longer than those associated with NCO events (Mitchel et al., 2010).

Studies of spontaneous or DNA damage-induced mitotic recombination in yeast have largely depended on selective systems that detect only one of the two CO products. An elegant system developed by Petes and colleagues allows both products of a CO event associated with LOH on chromosome IV or V to be captured within a sectored colony (Lee et al., 2009; St. Charles & Petes, 2013). Furthermore, if the diploid parent is derived from sequence-diverged haploids, GC tracts in each sector can be mapped using restriction-site polymorphisms or microarrays. Prior to the analysis of LOH-associated GC tracts, it was widely assumed that mitotic recombination would reflect the repair of a single broken chromatid. Analyses of sectored colonies suggested, however, that both chromatids were usually broken at `approximately the same position, with one break resolved as a NCO and the other as a CO. To explain the coincident breaks on sister chromatids, it was further proposed that the parent chromosome was broken prior to DNA replication, giving rise to two broken chromatids that were independently repaired (Lee et al., 2009). These studies thus changed the dogma concerning the timing of the damage that leads to LOH, while at the same time providing strong evidence that most spontaneous HR between homologs is initiated by DSBs, as has been widely assumed.

In the current study, we modified the chromosome IV mitotic LOH sectoring assay so that events were initiated by an I-*Sce*I-generated DSB. The repair-associated GC tracts in each sector were analyzed using microarrays, and results were consistent with the repair of two broken sister chromatids. Individual GC tracts within a given sector were then assigned to either the CO or NCO chromosome following their segregation into haploid meiotic products. Patterns of GC tracts in CO products suggested that ∼80% of hetDNA intermediates were repaired as GC events, with the remainder either escaping repair or reflecting restoration-type repair. In contrast to hetDNA tracts analyzed previously (Mitchel et al., 2010), there was no difference in the size or position of GC tracts associated with CO versus NCO events, with both frequently having GC tracts on both sides of the initiating DSB. This difference in hetDNA and GC tract locations derives from activity of the MMR machinery. The GC tracts characterized here are similar to those associated with spontaneous LOH events, and their properties are discussed in relation to current DSB-repair models.

## Results

A system developed in the Petes lab allows the detection of both products of reciprocal CO events on the right arm of chromosome IV as red-white sectored colonies (St. Charles & Petes, 2013). In this system, diploid strains are derived by crossing two sequence-diverged haploids (W303-1A and YJM789, hereafter abbreviated as W303 and YJM, respectively), allowing subsequent molecular characterization of the positions and lengths of GC tracts associated with individual reciprocal CO events. There are two genetic features of diploids relevant to CO detection: (1) homozygosity for the *ade2-1* ochre allele located on chromosome XV and (2) presence of a single, ectopic copy of the ochre-suppressing *SUP4-o* gene near the right end of one copy of chromosome IV. An *ade2-1/ade2-1* diploid produces a red pigment due to the accumulation of an intermediate in the adenine biosynthetic pathway, resulting in red colonies on adenine-limited medium. Complete suppression of the *ade2-1* alleles occurs when two copies of *SUP4-o* are present, and colonies are white instead of red. If only a single copy of *SUP4-o* is present, however, as in the starting diploid, there is partial suppression of the *ade2-1* alleles and colonies are pink.

As illustrated in Figure 2A, the YJM (black) homolog contains an ectopic *SUP4-o* allele inserted near the right end of chromosome IV (coordinate 1510386). When crossing over occurs after DNA replication, subsequent segregation of the CO chromosomes to opposite poles of the mitotic spindle produces a daughter cell with two copies of *SUP4-o* and a daughter with no copy of *SUP4-o.* In contrast to the starting diploid, which forms pink colonies, a CO that occurs at the time of plating gives rise to a red-white sectored colony. During DSB repair, information is transferred from the repair template to the broken chromosome to produce an associated GC tract. GC events are detected by monitoring the heterozygous versus homozygous status of single-nucleotide polymorphisms (SNPs) present in the W303 and YJM genomes (see below for further details). If only a single red chromatid is broken, repair gives rise to three chromatids with black information, which is referred to as a 3:1 conversion event (left side of Figure 2A). If both red chromatids are broken, however, each will receive information from the black repair template. The resulting gene conversion tracts on the repaired chromatids can be the same length or different lengths. In the former case, a 4:0 conversion tract is present in the sectored colony;; in the latter case, a hybrid 4:0/3:1 tract is produced (right side of Figure 2A). Spontaneous COs are usually associated with 4:0 or 4:0/3:1 hybrid tracts, suggesting that most DSBs are formed prior to S phase and are replicated to produce sister chromatids broken at the same position (Lee et al., 2009;; St. Charles & Petes, 2013).

**Figure 2.**
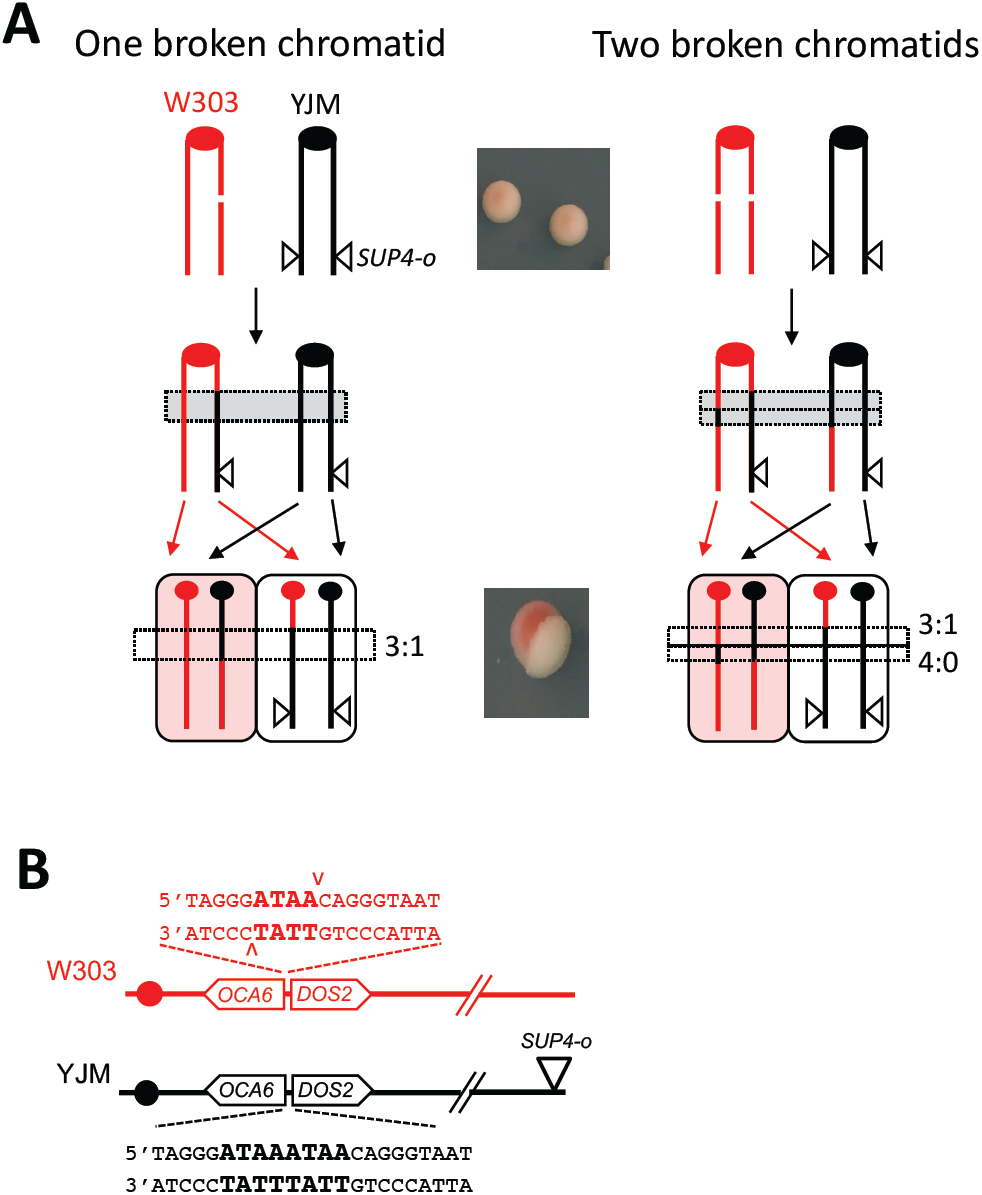
Detection of DSB-induced reciprocal COs. Red and black lines indicate replicated, sister chromatids of the W303 and YJM chromosome IV homologs, respectively;; filled circles corresponding to centromeres. **(A)** Red-white sectors correspond to reciprocal CO events. The starting diploid is pink because it contains a single copy of *SUP4-o* and suppression of the *ade2-1* alleles is incomplete. A CO followed by segregation of sister chromatids into different daughter cells yields a cell with no copy of *SUP4-o* and a cell with two copies. The former and latter produce the red and white sides of sectored colony. If only one of the W303/red sister chromatids is broken (left side), the transfer of information from a donor YJM/black chromatid produces a 3:1 GC tract. If both red chromatids are broken, a hybrid 4:0/3:1 tract is produced. **(B)** Sequence inserted into the W303 and YJM chromosomes. Only the W303 chromosome is cleaved by I-*Sce*I.

### Modifying the red-white sectoring system to initiate COs with a defined DSB

The red-white sectoring assay described above has been used extensively to characterize spontaneous and DNA-damage induced GC tracts associated with CO events in wild-type and mutant strains (Andersen et al., 2016; Lee et al., 2009; O'Connell, Jinks-Robertson, & Petes, 2015;; St. Charles & Petes, 2013;; Tang, Dominska, Gawel, Greenwell, & Petes, 2013;; Yin & Petes, 2014, 2015). With the exception of one study involving a trinucleotide-repeat recombination hotspot on chromosome V (Tang et al., 2013), however, the relationship between GC tracts and the position of the initiating lesion could not be determined. We thus modified the MMR-proficient W303/YJM hybrid diploid so that COs are initiated by a defined DSB, creating strain SJR4317 (Figure 2B). This was accomplished by inserting an 18-bp recognition site for the I-*Sce*I endonuclease on the right arm of the W303 homolog between the *OCA6* and *DOS2* loci (coordinate 583534);; this position is 133 kb distal to the centromere and 927 kb proximal to the *SUP4-o* marker. To induce a site-specific DSB, we inserted a galactose-inducible I-*Sce*I gene at the *HIS3* locus on one copy of chromosome XV. Finally, to eliminate the non-homology associated with insertion of the I-*Sce*I site into the W303 homolog, a non-cleavable I-*Sce*I site was inserted at the identical position on the YJM homolog. The non-cleavable site was created by duplicating the 4 bp flanked by I-*Sce*I generated nicks. Following repair of a DSB on a W303 chromatid using the YJM homolog as a template, the chromatid is refractory to further I-*Sce*I cleavage.

To introduce a DSB on the W303 homolog of SJR4317, we added 2% galactose to cells growing exponentially in rich medium containing 2% raffinose. After incubation for an additional 45 or 90 min, cells were plated on glucose-containing medium and red-white sectored colonies were scored microscopically. Without the addition of galactose, no red-white sectored colonies were detected among 8905 colonies analyzed. The frequency of red-white sectors was 2.7 x 10^-3^ (12/4464) and 6.3 x 10^-3^ (43/6855) after incubation with galactose for 45 and 90 min, respectively. Although we did not detect red-white sectored colonies in the absence of galactose, prior studies reported a frequency of 3.1 x 10^-5^ in a nearly isogenic strain (St. Charles & Petes, 2013). There was thus a robust increase in CO events, all of which analyzed (see below) were consistent with site-specific initiation by I-*Sce*I cleavage of the W303 chromosome.

### Microarray-based analysis of DSB-associated gene conversion tracts

The haploid parents of SJR4317 are diverged at the DNA sequence level and contain ∼55,000 SNPs. Approximately 1000 SNPs on the right arm of chromosome IV were monitored using a custom microarray containing oligonucleotides (oligos) specific for the W303 and YJM versions of each SNP (St. Charles & Petes, 2013). To determine the heterozygous versus homozygous status of SNPs in CO products, DNA from individual red or white sectors was labeled with a Cy5-tagged nucleotide and mixed with control, parental DNA labeled with a Cy3-tagged nucleotide. Following competitive hybridization of the mixed samples to the microarray, the ratio of Cy5 to Cy3 hybridization to each oligo was determined. A hybridization ratio of ∼1 to both W303- and YJM-specific oligos signals heterozygosity for the corresponding SNP. A hybridization ratio >1.5 for a haploid-specific oligo indicates homozygosity for the corresponding SNP, while a ratio <0.5 to the alternative haploid-specific oligo indicates loss of the corresponding SNP. Homozygosity of a SNP in only one side of a sectored colony corresponds to a 3:1 conversion event, while homozygosity in both sectors reflects a 4:0 conversion event. Because it is the red, W303 homolog that is cleaved by I-*Sce*I, conversion events are expected to involve unidirectional transfer of black, YJM-specific SNPs.

When both chromatids are broken, each half of a sectored colony contains a NCO chromosome as well as one of the CO chromosomes (see Figure 2B). In the current context, a NCO chromosome refers to one that maintains the parental linkage between the centromere and telomere, without regard to its participation in recombination or the presence/absence of GC tracts. An example of microarray results for the red and white halves of a single sectored colony (colony #2) is shown in Figure 3. There is reciprocal LOH that initiates near the induced DSB and extends to the end of the chromosomes in each sector (Figure 3A). Because SJR4317 is wild type with respect to the MMR pathway, SNP-containing hetDNA formed during DSB repair is expected to be repaired. In the red sector, this results in two transitions in addition to the transition at the site of the DSB (vertical dotted and solid black lines, respectively, in Figure 3B). In the white sector, however, there is only a single transition from SNP heterozygosity to homozygosity and this is centromere-proximal to the DSB. This is consistent with the presence of an unchanged, NCO YJM/black chromosome plus a CO chromosome containing the centromere of W303 in the white sector. Considering both sectors, three chromosomes contain YJM-specific, black SNPs to the left (centromere proximal) of the DSB and all four contain black SNPs immediately to the right of the break, which correspond to 3:1 and 4:0 GC tracts, respectively. As noted previously, a 4:0 tract indicates the cleavage and repair of both W303 chromatids. Finally, the 4:0 tract is followed by a second 3:1 GC tract before reciprocal LOH begins, consistent with the creation of GC tracts of different lengths during the independent repair of broken sister chromatids.

**Figure 3.**
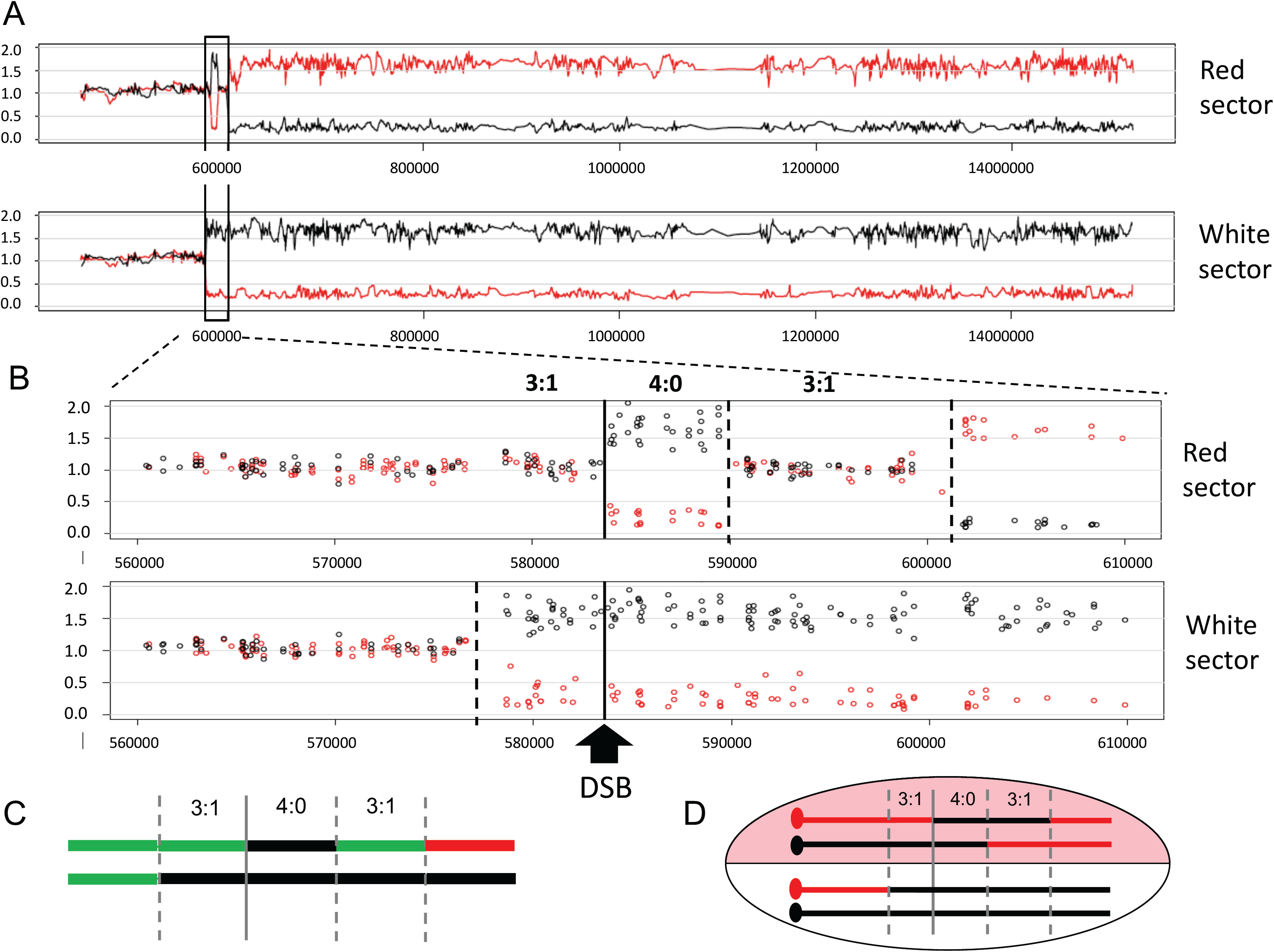
Microarray analysis of a red-white sectored colony. The data obtained with colony #2 are shown, along with the data interpretations. Black and red lines/circles correspond to W303- and YJM-specific SNPs, respectively. **(A)** Data from the right arm of chromosome IV are shown for the red and white sectors. The boxed area is enlarged in **(B)** to show the position of the DSB (black line) and transitions from heterozygosity to homozygosity (dotted black lines). 3:1 and 4:0 GC tracts are labeled. **(C)** A shorthand depiction of the chromosomes in the white and red sectors (top and bottom lines, respectively). The green lines correspond to heterozygosity. **(D)** The linkage relationships of the GC tracts with respect to the CO and NCO chromosomes in the red sector are illustrated. These were determined following the segregation of chromosomes into the haploid meiotic products.

A shorthand summary showing the heterozygous to homozygous transitions in each sector of a colony was developed by the Petes lab (St. Charles & Petes, 2013), and this style of presentation is presented in Figure 3C for the corresponding microarray data. In this depiction, black and red lines indicate homozygosity for YJM and W303 SNPs, respectively, while green lines correspond to heterozygosity. Twenty red-white sectored colonies were examined using SNP microarrays and the shorthand summaries of heterozygous to homozygous transitions in each are presented in Figure 4. For comparison purposes, transitions are aligned relative to the initiating DSB, and the summed length of all tracts within each colony is given. The summed GC tracts ranged from 7.3 to 32.3 kb, with a median length of 12.9 kb. With a single broken chromatid, the GC tract is always located on the CO chromatid and only a single transition between heterozygosity and homozygosity is expected in each sector (two transitions total). If both chromatids are broken, however, the GC can be on the NCO chromatid and this results in a total of three transitions between heterozygosity and homozygosity (see Figure S1). Of the 20 DSB-induced events analyzed here, 85% contained either a 4:0 tract (13/20) or a 3:1 GC pattern consistent with two broken chromatids (4/20; colonies #4, 6, 9 and 13); the remaining 15% (3/20; colonies #3, 11 and 12) had a 3:1 GC pattern consistent with breakage of a single chromatid. More detailed analysis of the chromosomes in the last class revealed that both W303 chromatids had lost the I-*Sce*I cleavage site, demonstrating that one chromatid had been repaired with no detectable acquisition of information from the donor (see below). Finally, in most sectored colonies (17/20) there was a 3:1 and/or 4:0 tract on each side of the initiating DSB.

**Figure 4.**
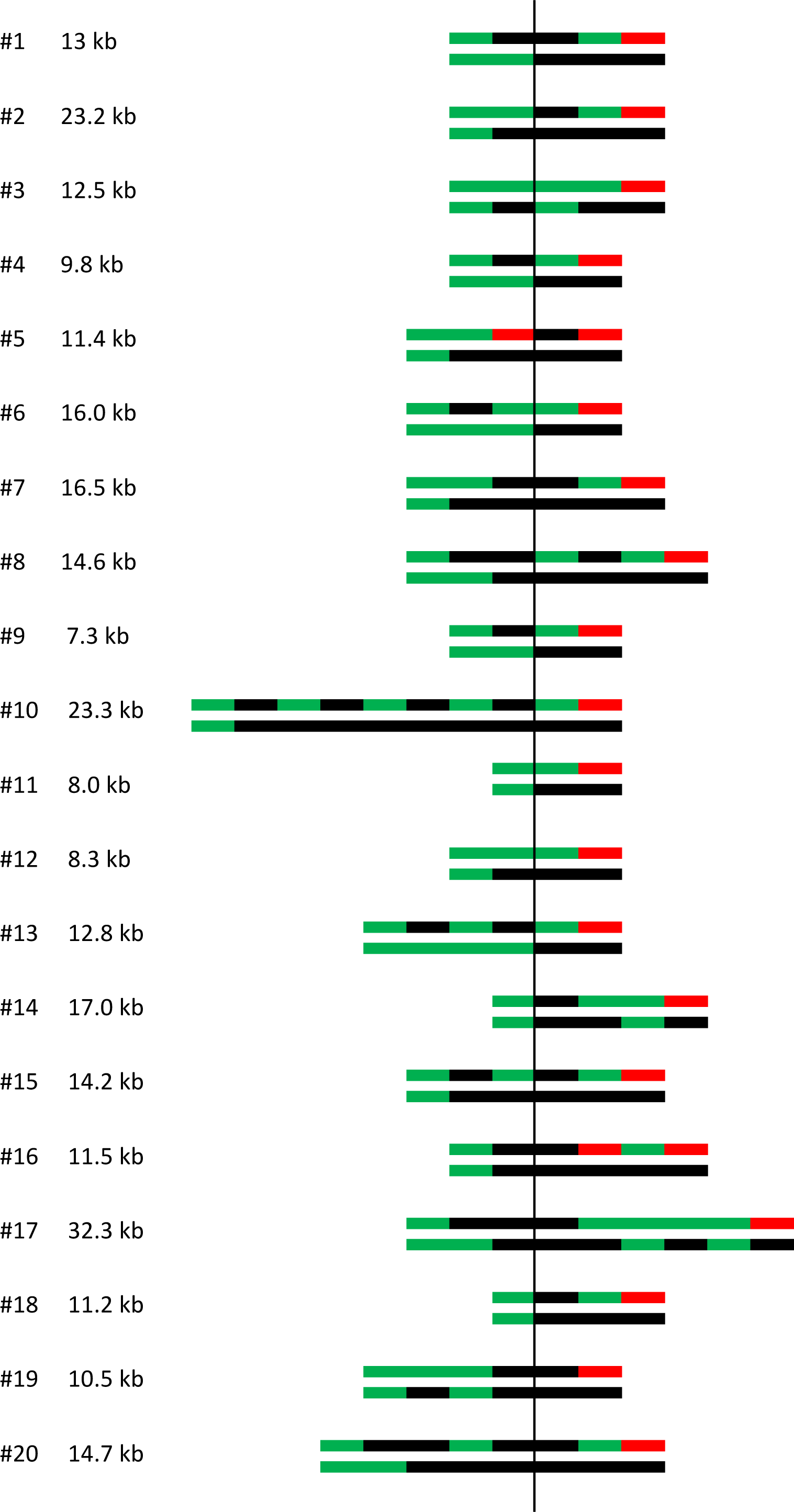
Summary of the transitions from heterozygosity to homozygosity in red-white sectored colonies. The shorthand depiction of chromosomes in each sector of the 20 colonies analyzed is presented as in Figure 3C. Transitions are aligned relative to initiating DSB (vertical line), and the GC tract length in each colony is given. The GC length was determined as the distance between the position of the first centromere-proximal transition from heterozygosity to homozygosity and the last, telomere-proximal transition from heterozygosity to terminal homozygosity. Multiple transitions most often occurred in the red sector, which contains a CO chromosome and the repaired, NCO red chromosome. Blocks of heterozygosity and homozygosity are not to scale.

### Assigning GC tracts to CO and NCO repair products

Hybrid 4:0 and 3:1 tracts are interpreted as two independent GC tracts of different lengths, but microarray data provided no information as to whether the 3:1 tract resides on the CO or the NCO chromosome. The locations of these and more complex GC tracts were obtained by sporulating and dissecting diploids from sectors that contained more than a single transition from heterozygosity to homozygosity. SNP linkages in haploid segregants were then determined using restriction site polymorphisms or by sequencing (see Material and Methods for details). In the red sector shown in Figure 3, for example, the GC tract that was part of the 4:0 event was 6.6 kb while that associated with the more extensive 3:1 event was 17 kb. Tetrad analysis revealed that the longer, 17 kb tract resided on the NCO rather than the CO chromosome; the positions of GC tracts on the four chromosomes in the sectored colony are summarized in Figure 3D. A similar depiction of the CO and NCO chromosomes in each sector of the 20 colonies analyzed is presented in Figure S2.

A repair scenario consistent with the microarray data in Figure 3 is presented in Figure 5. Following the cleavage of both red chromatids by I-*Sce*I (or cleavage in G1 and replication to produce two broken chromatids), the first repair event generates a dHJ that is resolved as a CO. Repair of the second DSB generates a NCO product and could, in principle occur by engaging any of the three, intact chromatids. In this particular case, the GC tract distal to the DSB on the recovered NCO chromosome is longer than that on the CO chromosome that contains the centromere of the YJM homolog, thereby eliminating the corresponding CO chromatid as the template. Because sister chromatids are preferred over homologous chromosomes, the sister chromatid (i.e., the CO chromatid with the same centromere) is assumed to be the most likely repair template. If repair of both broken sister chromatids occurs at the same time, however, the CO and NCO events would have to involve different chromatids of the homolog. It should be noted that GC tracts with a common endpoint (e.g., the non-hybrid, 4:0 tracts in colonies #5, 15, 16 and 19 in Figure S2) can arise if the second, NCO-type repair event uses the homolog-associated CO chromatid as the template.

**Figure 5.**
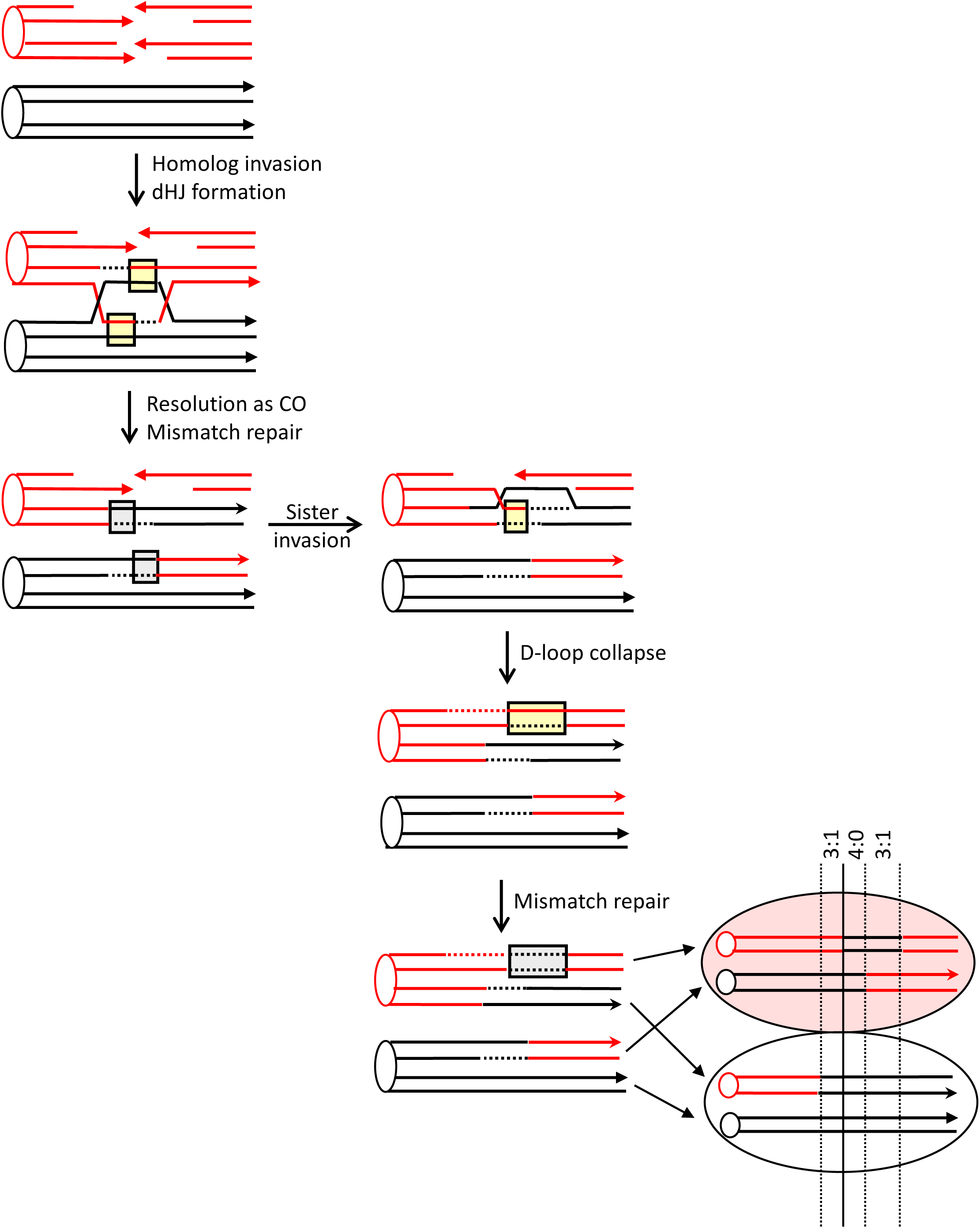
A DSB repair scenario consistent with the transitions in sectored colony #2. Red and black lines represent single strands of W303 and YJM chromosomes, respectively. Both red, sister chromatids are broken, and the first repair event is resolved as a CO. Each of the hetDNA tracts (yellow boxes) that flanks the DSB is repaired as a GC (gray boxes). The second repair event uses either the CO-resolved sister chromatid or the NCO chromatid of the homolog as a template. Repair using the sister chromatid is shown, and SDSA is assumed to be the repair mechanism because there is a GC tract on only one side of the initiating DSB. Because the hetDNA correction results in a GC tract that is longer than that on the CO chromosome, the non-sister, CO chromatid is excluded as a repair template. For this particular sectored colony, the NCO event could have preceded that of the CO event or both could have occurred at the same time. In this case, both repair events would have involved the homolog.

### Gene conversion tracts in CO products

CO-associated gene conversion (and restoration) tracts in the 20 sectored colonies are presented in Figure 6A, where the CO chromosomes within each colony are paired. Colonies are arranged based on the total extent of strand transfer detected, which ranged from none to almost 25 kb;; only one CO (colony #4) lacked an associated GC tract. The first SNPs on the left and right sides of the DSB were 114 and 218 bp, respectively from the corresponding 3′ ends created by I-*Sce*I cleavage. For all CO pairs, the transition from red to black information was consistent with strand transfer initiating at the DSB. When GC tracts that spanned the DSB were considered as a single tract, the median tract length was 10.8 kb. When the lengths of GC tracts on opposing sides of the DSB were considered separately, those on the left (centromere-proximal) and right sides of break were similar: 6.1 kb and 7.4 kb, respectively.

**Figure 6.**
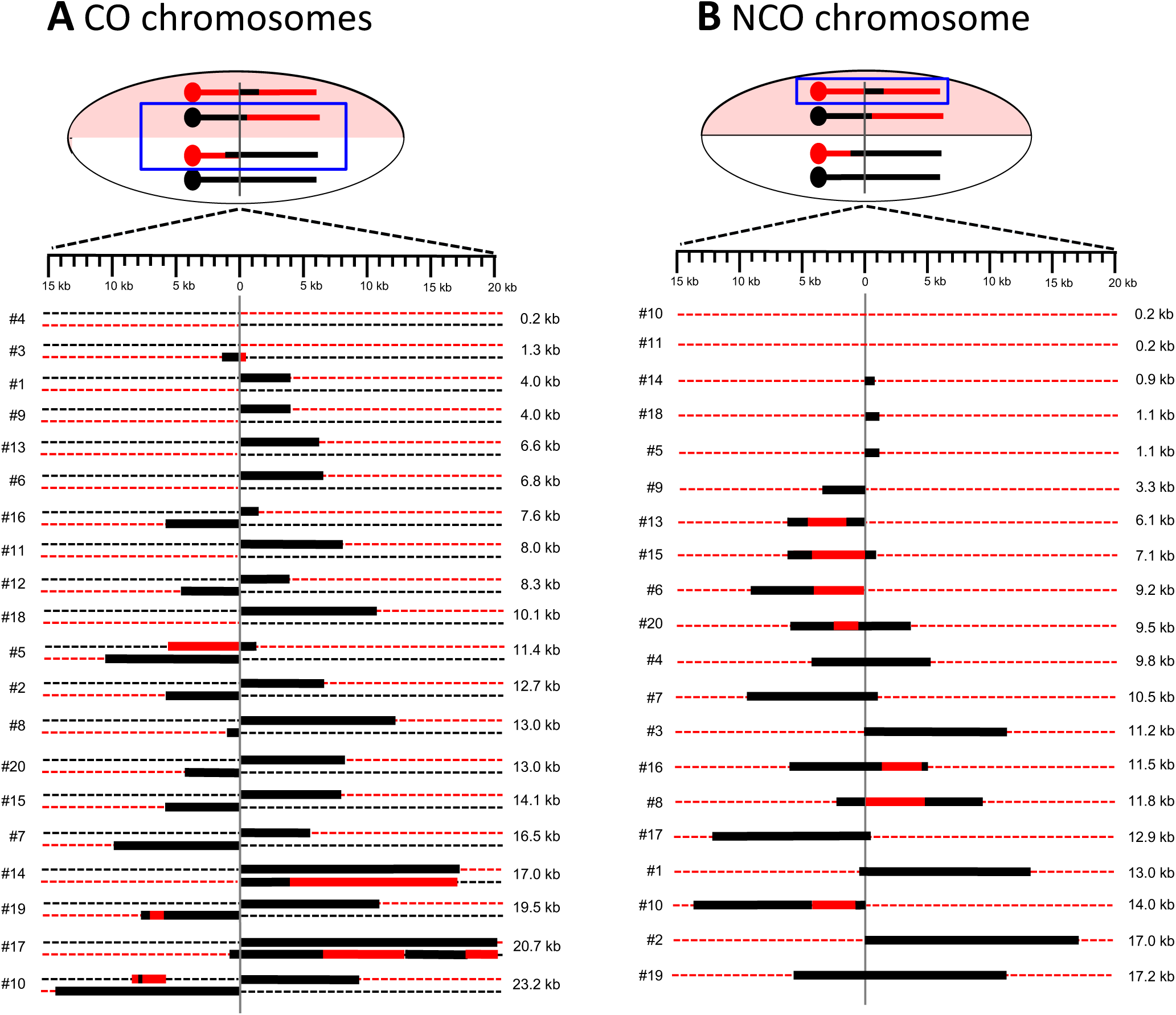
GC tracts associated with CO and NCO repair products. Dotted red and black lines correspond to W303 and YJM chromosomes, respectively. Thick black and red lines are GC and restoration tracts, respectively, and are shown to scale. The vertical gray line indicates the position of the initiating DSB. Events are arranged based on the total GC tract length, from smallest to largest. **(A)** CO chromosomes present in each sectored colony. The total GC length was obtained by summing the lengths on each side of the DSB;; symmetric tracts were considered a single tract for the length calculation. **(B)** GC tracts associated with the NCO event in each sectored colony are shown. There were no changes to the black/YJM NCO chromosome, so only the repaired red/W303 chromosome is shown.

In our previous analyses of CO-associated hetDNA in a plasmid-based assay, there was always a hetDNA tract on each side of the initiating DSB, as predicted by the classic DSB repair model (Mitchel et al., 2010). In the current study, however, 40% of CO events (8/19) had a GC tract that was confined to only one side of the DSB. One explanation for the asymmetry observed here is that hetDNA tracts are often very short (only 100-200 bp) on one side of the initiating break and, therefore, undetectable. Such asymmetry was not evident in the CO products that had a GC tract on both sides of the DSB or in our previous analysis of CO-associated hetDNA (Mitchel et al., 2010), but has been noted in meiotic studies (Jessop, Allers, & Lichten, 2005; Merker, Dominska, & Petes, 2003). We suggest that the difference between the locations of mitotic hetDNA and GC tracts may reflect the mismatch correction process.

MMR-directed removal of the strand contributed by the broken molecule results in GC, while removal of the strand contributed by the repair template yields a genetically silent restoration event. Assuming that the 19 COs with detectable GC tracts initially had hetDNA that covered at least one SNP on each side of the initiating break (38 hetDNA tracts total), there were 30 GC and 8 restoration events among the events analyzed. hetDNA repair is thus biased, but not completely so, during mitotic recombination and generates a GC event ∼80% of the time. An alternative explanation is that mismatch correction frequently failed to occur, with subsequent segregation of unrepaired hetDNA yielding apparent restoration as well as GC events. It is interesting to note that among the eight COs with a GC tract on only one side of the DSB, seven had the tract on the right side of the break (p=0.08). Although the number of events analyzed here was small, the asymmetry in GC position could reflect an invading-end bias coupled with biased repair of mismatches on the invading end versus the end captured by the D-loop.

The GC tracts in most sectored colonies were consistent with continuous repair of hetDNA intermediates, but there also were discontinuous tracts. In colony #19, for example, the GC tract on the left side of the DSB was interrupted by a tract of apparent restoration. One explanation for such discontinuities is the occurrence of discontinuous/patchy MMR tracts. As noted above, restoration could reflect either repair that removes the donor information or the complete absence of repair, with subsequent segregation of donor and recipient strands. Alternatively, it is possible that frequent template switching occurs during DSB repair. According to this model, the discontinuous pattern in CO #19 would be produced if the black homolog were initially invaded and used to prime DNA synthesis. Following dismantling of the D-loop, the red sister chromatid would then be invaded by the displaced, extended strand and used as a template for additional DNA synthesis. A final template switch back to the black template would then give rise to the break-distal GC tract. In addition to discontinuous GC tracts, there were four colonies that either contained GC tracts at the same position in both CO products or had a tract of restoration opposite a GC tract (#5, #10, #14 and #17). We suggest that these events reflect branch migration of an HJ away from the site of the DSB, converting asymmetric hetDNA that is localized to only one of the interacting duplexes to symmetric hetDNA that covers the same SNPs on both duplexes. Both patchy MMR and branch migration have been previously invoked to explain complex GC patterns among spontaneous CO events (St. Charles & Petes, 2013).

### Gene conversion tracts in NCO products

Figure 6B summarizes the positions and lengths of GC tracts associated with the DSB-induced NCO events. Within all colonies examined, the black/YJM NCO chromosome, which lacked the I-*Sce*I cut site, was unaltered;; only the red/W303 NCO chromosome is shown. The W303 NCO chromosome contained a GC tract in all but two of the sectored colonies (#10 and 11). The absence of a GC tract could reflect an uncut chromatid, a very small amount of strand exchange (less than ∼200 bp), repair of the hetDNA intermediate as a restoration rather than a GC event, or segregation of unrepaired hetDNA. To determine whether the parental chromatids had been broken and repaired without a residual GC footprint, we examined each for presence/absence of the I-*Sce*I cleavage site. In both cases, the I-*Sce*I site was absent. Thus, in all sectored colonies analyzed here, both sister chromatids were broken and repaired.

Half (9/18) of the NCO chromosomes had a GC tract on both sides of the initiating DSB (a bidirectional tract), which is consistent with either dissolution of a double HJ or a double-SDSA event. The other half contained a unidirectional GC tract, which is the pattern predicted by the SDSA pathway (see Figure 1). The median length of the unidirectional GC tracts was 6.1 kb, while that of bidirectional GC tracts was 11.6 kb. As seen for CO-associated GC tracts, NCO-associated tracts were often patchy or discontinuous, with a tract of black, YJM-specific SNPs interrupted by a tract of red SNPs. Such discontinuous tracts were present in 40% (7/18) of repaired NCO chromosomes.

The equality between unidirectional and bidirectional GC tracts in the NCO events analyzed here is in striking contrast to the strong bias for unidirectional tracts reported previously in an ectopic system when hetDNA was monitored in the absence of the Mlh1 MMR protein (Guo et al., 2017). There are three differences between the system used here to examine GC tracts and that used previously to characterize hetDNA tracts. First, GC tracts were examined in an MMR-proficient background while hetDNA analysis requires that experiments be done in the absence of MMR. Second, GC tracts were examined between allelic sequences with unlimited homology, whereas the ectopic assay involved 4-kb repeats (Guo et al., 2017). Finally, GC tracts were examined in a diploid background while haploid strains were used to analyze hetDNA tracts. The potential contribution of MMR to the different positions of GC versus hetDNA tracts was addressed by performing the ectopic experiments in an MMR-proficient background. The ectopic system is shown in Figure 7A, and events were similarly initiated by I-*Sce*I cleavage. Whereas only 14/108 hetDNA tracts in the ectopic system were bidirectional in the absence of *MLH1*, 44/119 GC tracts were bidirectional in MMR-proficient cells (p<0.001 by Chi-square).

**Figure 7.**
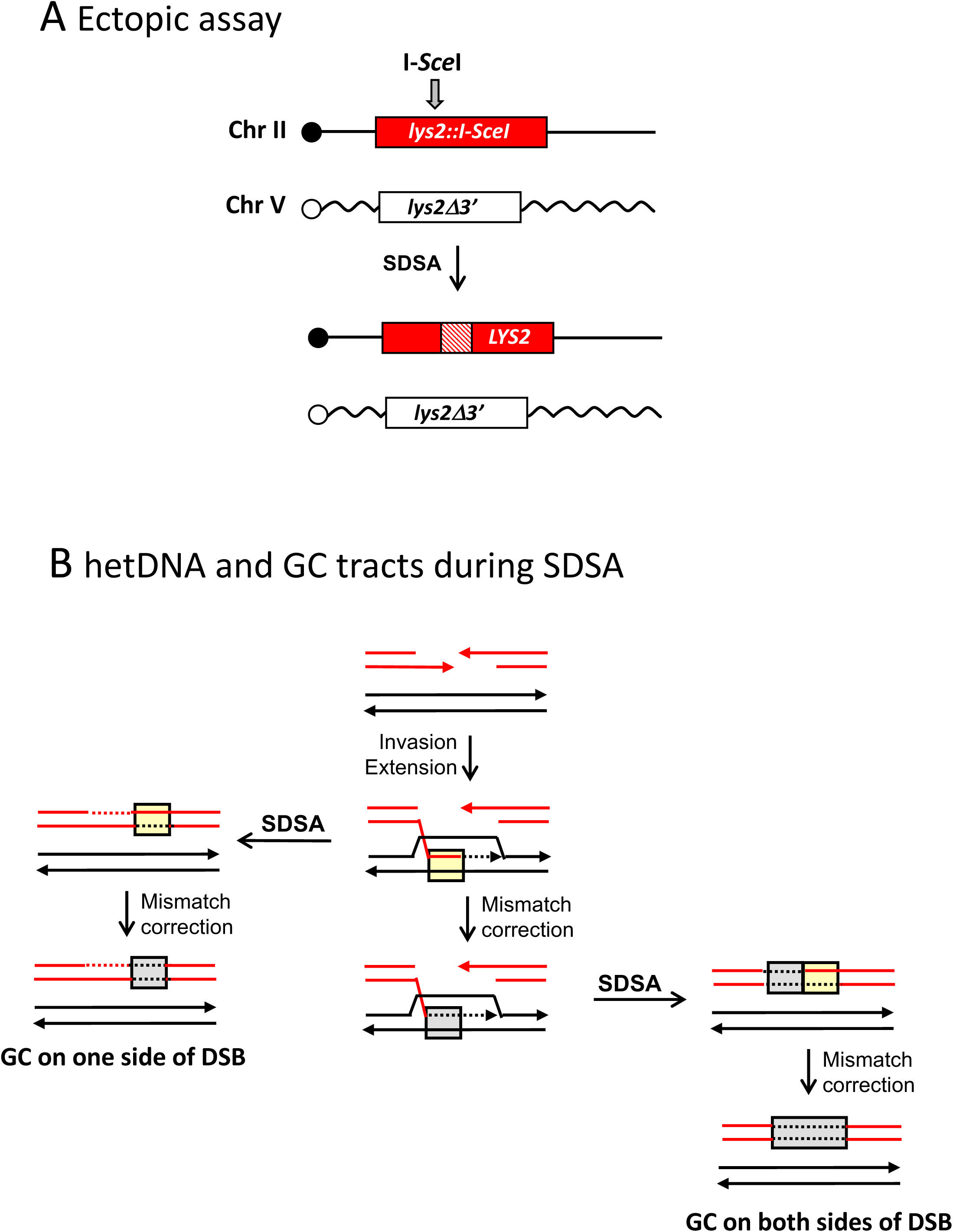
Mismatch repair during SDSA. **(A)** The ectopic assay uses diverged, 4.3 kb *lys2* repeats on chromosomes II and V. The full-length, recipient allele contains an intact I-*Sce*I cleavage site and the truncated, donor allele contains a mutated site;; the I-*Sce*I gene was fused to *pGAL* and integrated at *HIS3*. Following galactose induction of I-*Sce*I, repair of the broken allele produces a selectable, Lys^+^ phenotype. SDSA is inferred when the repaired allele contains a hetDNA tract (cross-hatched region) on only one side of the DSB and the donor allele is unchanged. hetDNA-and GC-tract positions were determined by sequencing products obtained in an MMR-defective (*mlh1*Δ) or wild-type background, respectively. **(B)** Products predicted from SDSA-mediated repair are illustrated. Strands of the broken, recipient allele are red and those of the donor allele are black. hDNA is in yellow boxes and GC tracts resulting from hetDNA repair are in gray boxes. If MMR occurs only after repair is complete, GC will be confined to one side of the initiating DSB (left side). If MMR can occur before the D-loop is dismantled, however, then there will a GC tract that spans the DSB (right side).

In the ectopic assay, there are two possible explanations for the shift from unidirectional hetDNA tracts in the absence of MMR to bidirectional GC tracts in the presence of MMR. First, it is possible that the MMR system alters the underlying mechanism of DSB repair. In this scenario, there would be an MMR-dependent shift from single-end engagement of the donor (unidirectional hetDNA tracts) to engagement of the donor by both DNA ends (bidirectional GC tracts). Engagement by both DNA ends would lead to formation of a fully-ligated double HJ-containing intermediate resolved by dissolution, or to a double-SDSA event. An alternative explanation is that the relative contribution of SDSA to NCOs remains the same in the presence and absence of MMR, but that the mismatches created by strand invasion are repaired prior to the dismantling of the D-loop intermediate (Figure 7B). The footprint of hetDNA created by strand invasion would thus persist as a GC event after D-loop collapse, rather than being permanently erased. With regard to the current analysis, we note that loss of Mlh1 prevents only the processing of mismatches;; mismatches within hetDNA will still be recognized and bound by MutS-like complexes.

## Discussion

The genetic system used here detects reciprocal LOH for a *SUP4-o* marker near the right end of chromosome IV, which results in a red-white sectored colony. Prior analyses of GC tracts associated with spontaneous LOH events were consistent with DSB-initiated recombination, and the prevalence of 4:0 GC tracts indicated that most breaks occurred in the G1 phase of the cell cycle (St. Charles & Petes, 2013). Replication of a broken chromosome yields sister chromatids broken at the same position, with one sister repaired as a CO event and the other as a NCO. Limitations associated with the analysis of spontaneous events were that, with one exception (Tang et al., 2013), the positions of the initiating lesions within GC tracts were unknown, and the basic parameters of GC tracts represented a compilation of events initiated at multiple sites. In addition, GC tracts within a given colony were considered jointly rather than as tracts associated with independent CO-and NCO-type repair events. To examine GC tracts initiated by a site-specific DSB, we inserted an I-*Sce*I cleavage site into one of the chromosome IV homologs used in the spontaneous LOH assay. Twenty sectored colonies isolated following I-*Sce*I induction were analyzed by SNP microarrays, and each reflected the repair of two broken chromosomes or chromatids: one repaired as a CO event and the other as a NCO. Segregation of the CO and NCO chromosomes into meiotic products allowed the GC tract(s) associated with each repair type to be analyzed separately as well as jointly. In the discussion that follows, we compare GC tracts associated with DSB-induced LOH events to those previously described for spontaneous LOH events. In addition, the GC tracts associated with NCO-versus CO-type repair events that initiate at a common site are compared and are discussed in relation to current models of DSB repair.

Following the introduction of a targeted DSB, the red-white sectoring frequency was approximately 100-fold greater than the spontaneous frequency, which reflects random initiation across an ∼1Mb region. The positions of GC tracts in all red-white colonies analyzed were consistent with initiation by I-*Sce*I cleavage and in all cases, both chromosomes or chromatids were broken. Our analyses cannot distinguish chromosome cleavage in G1, followed by replication of the broken chromosome, from independent cleavage of sister chromatids in G2. In the prior study of spontaneous LOH events on chromosome IV, there were four GC patterns consistent with single-chromatid breakage in G2 (St. Charles & Petes, 2013), and it is noteworthy that two of these G2 patterns were detected among the DSB-induced events examined here. The observance of a G2 initiation pattern in a context where we know that both chromosomes/chromatids were broken suggests that many, if not most, of the ∼25% of spontaneous events attributed to initiation in G2 may also have been G1-initiated events. Given that the sister chromatid is strongly preferred over the homolog as the repair template (Kadyk & Hartwell, 1992), G2-initiated events may rarely, if ever, give rise to LOH.

The total GC tract length within the sectored colonies, which corresponds to the region spanned when CO and NCO products are jointly considered, ranged from 7.3 to 32.3 kb, with a median length of 12.9 kb (Figure 4). This length is very similar to the previous estimate of 14.8 kb for spontaneous, G1-initiated GC tracts (St. Charles & Petes, 2013). For 90 spontaneous events with a GC pattern consistent with initiation in G1, 7 contained a single 4:0 tract, 45 had a simple hybrid 4:0/3:1 or 3:1/4:0/3:1 tract, and 38 had a more complex GC pattern. Although the sample size of DSB-induced events is much smaller in the current analysis, 14/17 DSB-associated tracts with the G1-initiation pattern were classified as complex. This proportion is much larger than reported for spontaneous events (p=0.003) and may partially reflect the very short distance between the targeted DSB and flanking SNP heterozygosities (114 and 218 bp, respectively, on the left and right sides of the DSB). Not surprisingly, the perceived complexity was reduced when the CO and NCO events within a given sectored colony were analyzed separately (Figure 6).

Current models of DSB repair make specific predictions about where hetDNA should be located relative to the initiating break (see Figure 1). We previously mapped hetDNA in the NCO and CO products obtained following repair of a linearized plasmid (Mitchel et al., 2010), as well as in DSB-induced NCO events generated in an ectopic, chromosomal assay (Guo et al., 2017). In both systems, experiments were done in the absence of the Mlh1 MMR protein in order to preserve mismatches that define the extent/position of hetDNA. hetDNA tracts associated with CO events in the plasmid-based assay were located on each side of the initiating break (bidirectional tracts) as predicted by the classic model of DSB repair. Each CO product contained hetDNA on one side of the initiating DSB, and the hetDNA tracts in the CO products were on opposite sides of the break (Mitchel et al., 2010). In an MMR-proficient background, the repair of hetDNA is expected to produce a bidirectional GC tract that spans the DSB in the reciprocal repair products (Figure 1), and this was observed in the plasmid-based assay. Among the COs examined here, however, only 11/20 contained a GC tract that spanned the break, and one event contained no associated GC tract (Figure 6A). We suggest that the reduction in CO-associated, bidirectional GC tracts in the allelic assay relative to bidirectional hetDNA tracts observed in the plasmid assay reflects either restoration-type repair and/or the segregation of unrepaired hetDNA. If solely the result of repair, then the estimated frequency of restoration is ∼20%.

For NCO events examined previously in the plasmid and ectopic assays, all hetDNA was confined to the broken molecule, suggesting little, if any production of NCOs via HJ cleavage (Guo et al., 2017;; Mitchel et al., 2010). Furthermore, 80-90% of hetDNA in the repaired allele was confined to a single side of the initiating DSB, consistent with most repair occurring via the SDSA pathway. When the MMR machinery is intact, NCOs that arise via the SDSA pathway are expected to have a GC tract on only one side of the initiating DSB, and this was observed in a plasmid-based assay (Mitchel et al., 2010). Although NCO events produced by HJ cleavage or dissolution are associated with distinctive hetDNA positions, they are expected to be identical in terms of GC tracts. GC tracts are thus generally less informative with regard to the underlying molecular mechanism than are hetDNA tracts (see Figure 1).

Based on our prior analysis of hetDNA and GC tracts in a plasmid-based assay (Mitchel et al., 2010), we anticipated that most allelic NCOs in the LOH assay would reflect SDSA and hence would have a GC tract on only one side of the initiating DSB. Among the NCO events with GC tracts, however, 50% (9/18) had a GC tract on each side of the DSB (Figure 6B). If one assumes that SDSA is the predominant NCO mechanism, one explanation for the frequent bidirectional GC tracts is correction of hetDNA created by strand invasion prior to disassembly of the D-loop intermediate (Figure 7B). In the absence of MMR the footprint of the hetDNA created by strand invasion will be permanently lost upon D-loop collapse. To examine this possibility, we determined the positions of DSB-induced hetDNA and GC tracts in an ectopic assay by performing experiments in absence and presence of Mlh1, respectively. Whereas only ∼10% of hetDNA tracts were bidirectional in the absence of MMR, ∼30% of the GC tracts in the ectopic assay were bidirectional in the presence of MMR. This shift is consistent with the MMR-dependent preservation of a transient hetDNA footprint that is formed on the invading side of a DSB during SDSA. Despite the differences in the ectopic and allelic assays, the proportion of bidirectional GC tracts in the ectopic (44/119) and allelic (9/18) systems were similar (p=0.31 by Fisher Exact) when experiments were done a MMR-proficient background. An alternative explanation for more bidirectional events in the presence than in the absence of MMR is that the MMR system alters the mechanism of DSB repair. The reason for the agreement between hetDNA and GC tract positions in the plasmid-based assay (Mitchel et al., 2010) is unclear, but may be related to the very small size of substrate homology (∼800 bp). The stability of the initiating D-loop is likely related to the extent of hetDNA formed upon strand invasion, and this is limited in the plasmid assay. A more unstable D-loop may, in turn, be dismantled before hetDNA repair can occur.

In the LOH assay it should be noted that the proportions of gene conversion tracts on one versus both sides of the initiating DSB were indistinguishable for NCO and CO events (Figure 6). In addition, the median sizes of GC tracts associated with CO and NCO events were the same (10.8 and 9.7 kb, respectively). These similarities may be coincidental, but also are suggestive of a common mechanism/intermediate. This raises the possibility that NCOs as well as COs are generated via HJ cleavage in the LOH system. It is possible, for example, that the SDSA mechanism only becomes a significant contributor to NCOs when homology between the recombination substrates is limited, as in an ectopic or plasmid-based assay. This could occur, for example, if D-loop stability is inversely related to the length of hetDNA that forms during the initiating strand invasion event. hetDNA formation will be limited by the extent of homology present in the plasmid-based and ectopic assays, whereas there is no limitation in the allelic, LOH assay. Indeed, in the ectopic assay where NCO-associated hetDNA was examined, the available homology was 4.3 kb (Guo et al., 2017) which is only one-half the median length of NCO-associated GC tracts in the LOH assay. Determining whether the mechanism of NCO production is related to the extent of available homology will require an examination of hetDNA relative to the initiating break in the LOH assay.

For both the CO and NCO events analyzed in the LOH assay, a significant fraction of events was more complex than predicted by current DSB repair models. This complexity included the interspersion of restoration/unrepaired tracts and GC tracts (∼25% of CO and NCO events) and the occurrence of symmetric GC/restoration tracts (∼20% of CO events). The former could reflect patchy MMR and/or template switching. A recent analysis of hetDNA associated with spontaneous and UV-induced LOH events reached similar conclusions in terms of the complexity of recombination intermediates (see the accompanying paper by Yin and Petes). Additionally, hetDNA complexity has been observed in studies of meiotic recombination (Martini et al., 2011). Overall, however, the molecular analyses of mitotic and meiotic products of recombination are generally consistent with current DSB repair models. Given the high conservation of recombination mechanisms, the results reported here are likely relevant to LOH in mammalian cells, which is an important contributor to tumor-supporessor loss and cancer development.

## Acknowledgments

The authors are grateful to Tom Petes for helpful discussions throughout this work, and to members of his lab for assistance in the microarray analyses. Work in the SJR lab is supported by NIH grant R35 GM118077; YFH was partially supported by a pre-doctoral grant from the American Heart Association.

## Materials and Methods

### Yeast strain constructions

All experiments were done in a diploid SJR4317, which was constructed by crossing derivatives of the diverged haploids W303-1A (Thomas & Rothstein, 1989) and YJM789 (Wei et al., 2007). A complete list of intermediate strains is provided in Table S1. Strains JSC12-1 and JSC21-1 were the starting W303-1A and YJM746 derivatives, respectively, and were described previously (St. Charles and Petes 2013). The relevant feature of these strains is the insertion of the *SUP4-o* and *kanMX* markers at allelic positions near the right end of chromosome IV (coordinate IV1510386 in the *Saccharomyces* Genome Database). Either a cleavable or non-cleavable I-*Sce*I recognition site was inserted centromere-proximal to these markers using the two-step *delitto perfetto* method (Storici and Resnick 2003). First, a PCR-generated CORE-UH cassette was introduced at corrdinate IV583526 between the *OCA6* and *DOS2* genes. The CORE-UH cassette was then replaced with a PCR-generated fragment containing a cleavable, 18 bp I-*Sce*I recognition site (5’-TAGGGATAACAGGGTAAT) or a 22 bp, non-cleavable site (I-*Sce*Inc; 5’-TAGGGATAAATAACAGGGTAAT). The I-*Sce*I and I-*Sce*Inc sites were inserted into the W303 and YJM homologs, respectively, and the resulting haploids were crossed to generate a W303/YJM diploid strain. Finally, SJR4317 was derived by inserting a galactose-inducible I-*Sce*I gene amplified from pGSHU (Storici 2003) at the *HIS3* locus of the W303 chromosome XV homolog, and deleting the *MAT*α locus on from YJM chromosome III homolog in order to prevent unwanted sporulation.

### I-*Sce*I induction and screening for DSB-induced COs

Prior to I-*Sce*I induction, yeast cells were grown overnight at 30°C in 5 ml of non-selective medium containing 1% yeast extract, 2% bacto-peptone and 2% raffinose (YEPR). Raffinose is a neutral carbon source that neither induces nor suppresses the galactose promoter. Single colonies were used to inoculate YEPR cultures, and following overnight growth, cells were then diluted to an OD of 0.4 in fresh YEPR medium and split into two parallel cultures. When cells reached an OD of 0.8-1, galactose (2% final concentration) was added to one culture of each pair and cells were incubated for an additional 45 or 90 minutes. Following induction for the specified time, yeast cells were pelleted and resuspended in water.

Approximately 500 cells/plate were spread on synthetic dextrose medium supplemented with uracil and all amino acids except arginine. Adenine was additionally present at 10 μg/ml, a limiting amount that allows red color development of Ade^-^ cells. Plates were incubated at 30°C overnight and at room temperature for additional 2 days before being moved to 4°C to allow red pigment development. Red-white sectored colonies were identified microscopically, and the sectoring frequency was calculated by dividing the number of sectored colonies by the total number of colonies screened. The frequencies of sectored colonies in non-induced cultures and in cultures induced for 90 min were based on data from three independent experiments; for the 45 min induction, data were from two independent experiments.

### Microarray analysis

Hybridization to custom, chromosome IV SNP microarrays and subsequent data analysis was as previously described (St. Charles and Petes 2013). Briefly, cells in each half of a sectored colony were purified by streaking on YEP-glucose medium, and a single colony from each was used for DNA isolation. Genomic DNA was extracted in agarose plugs and then sheared to 200-400 bp by sonication. DNA from each sector was labeled with Cy5-dUTP and mixed with control parental DNA labeled with Cy3-dUTP. Mixed DNA was competitively hybridized to a SNP microarray, and the ratio of Cy5 to Cy3 hybridization to each SNP was determined. Each SNP was represented by four 25-nt oligonucleotides that corresponded to the Watson and Crick strands of each homolog.

### Determining the positions of gene conversion tracts

Each sector contained a CO and NCO chromosome. When there was a single transition from heterozygosity to homozygosity, the transition was assigned to the CO chromosome; this was the case in almost all white sectors. In most red sectors (and some white sectors), however, there were multiple transitions and assigning GC tracts to the CO or NCO chromosome required additional analysis. For colony #1, for example, the red sector contained a region of GC (black homozygosity) that spanned the DSB. Going towards telomere, this was followed by region of green heterozygosity before a terminal region of red homozygosity (Figure 4). The region of centromere-distal heterozygosity reflects different CG tract lengths on the CO and NCO chromosomes, but provides no information as to which chromosome contains the longer tract. To determine this, the CO and NCO chromosomes were segregated into haploid colonies by sporulation and tetrad dissection, and were distinguished using a W303-YJM restriction site polymorphism in the centromere-proximal region of heterozygosity where there had been no exchange of genetic information. The genetic linkage of this polymorphism to one within the break-distal region of heterozygosity was then assessed. The primers used to amplify relevant segments and the corresponding restriction site polymorphisms are in Table S2 and Table S3 gives the primers used to analyze each sector. Primer positions relative to the initiating DSB are shown in Figure S3. For analysis of colony #1, for example, primer pair 6 amplified a centromere-proximal fragment that contained an *Afl*II cleavage site specific to YJM, and primer pair 11 amplified a fragment containing a *Nar*I site also specific for YJM. In dissected tetrads, these were linked as in the parent chromosomes in only 2/51 spores analyzed, which places the longer tract on the NCO chromosome. Spore data are summarized in Table S3.

## Supplemental Figure Legends

**Table S1:** Yeast strains.

| Strain | Relevant genotype | Comments/reference |
| --- | --- | --- |
| JSC12-1<br>(W3031A) | *MATa RAD5 ade2-1 leu2-3,112 his3-11,15 ura3-1 trp1-1*<br>*can1-100Δ::nat IV1510386::kanMX-can1-100* | Isogenic derivative of W303-1A |
| JSC21-1<br>(YJM789) | *MATα ade2-1 ura3 gal2 ho:hisG canΔ::NAT*<br>*IV1510386::SUP4-o* | Isogenic derivative of YJM789 |
| SJR4246<br>(W303) | *IV1510386::kanMX IV583526:CORE-UH* | CORE-UH cassette inserted between the OCA6 and *DOS2* loci of JSC12-1 |
| SJR4248<br>(YJM) | *IV1510386::SUP4-o IV583526:CORE-UH* | CORE-UH cassette inserted between the OCA6 and *DOS2* loci of JSC21-1 |
| SJR4250<br>(W303) | *IV1510386::kanMX IV583534:I-SceI* | CORE-UH of SJR4246 replaced with 18 bp I-SceI cut site |
| SJR4252<br>(YJM) | *IV1510386::SUP4-o IV583534:I-SceI<sup>nc</sup>* | CORE-UH of SJR4248 replaced with 22 bp mutant (non-cutting, nc) I-SceI site |
| SJR4313<br>(W303/YJM) | *MATa/MATα IV1510386::kanMX/SUP4-o IV583534:I-SceI/I-SceI<sup>nc</sup>* | SJR4250 x SJR 4252 |
| SJR4315<br>(W303/YJM) | *MATa/matαΔ::URA3 IV1510386::kanMX/SUP4-o*<br>*IV583534:I-SceI/I-SceI<sup>nc</sup>* | *MATα* locus of SJR4313 replaced with *URA3* to prevent sporulation. |
| SJR4317<br>(W303/YJM) | *MATa/matαΔ::URA3 IV1510386::kanMX/SUP4-o*<br>*IV583534:I-SceI/I-SceI<sup>nc</sup> his3Δ::pGAL-I-SceI-hph/HIS3* | *his3-11,15* replaced with a *pGAL-I-SceI* fusion |

**Table S2.**
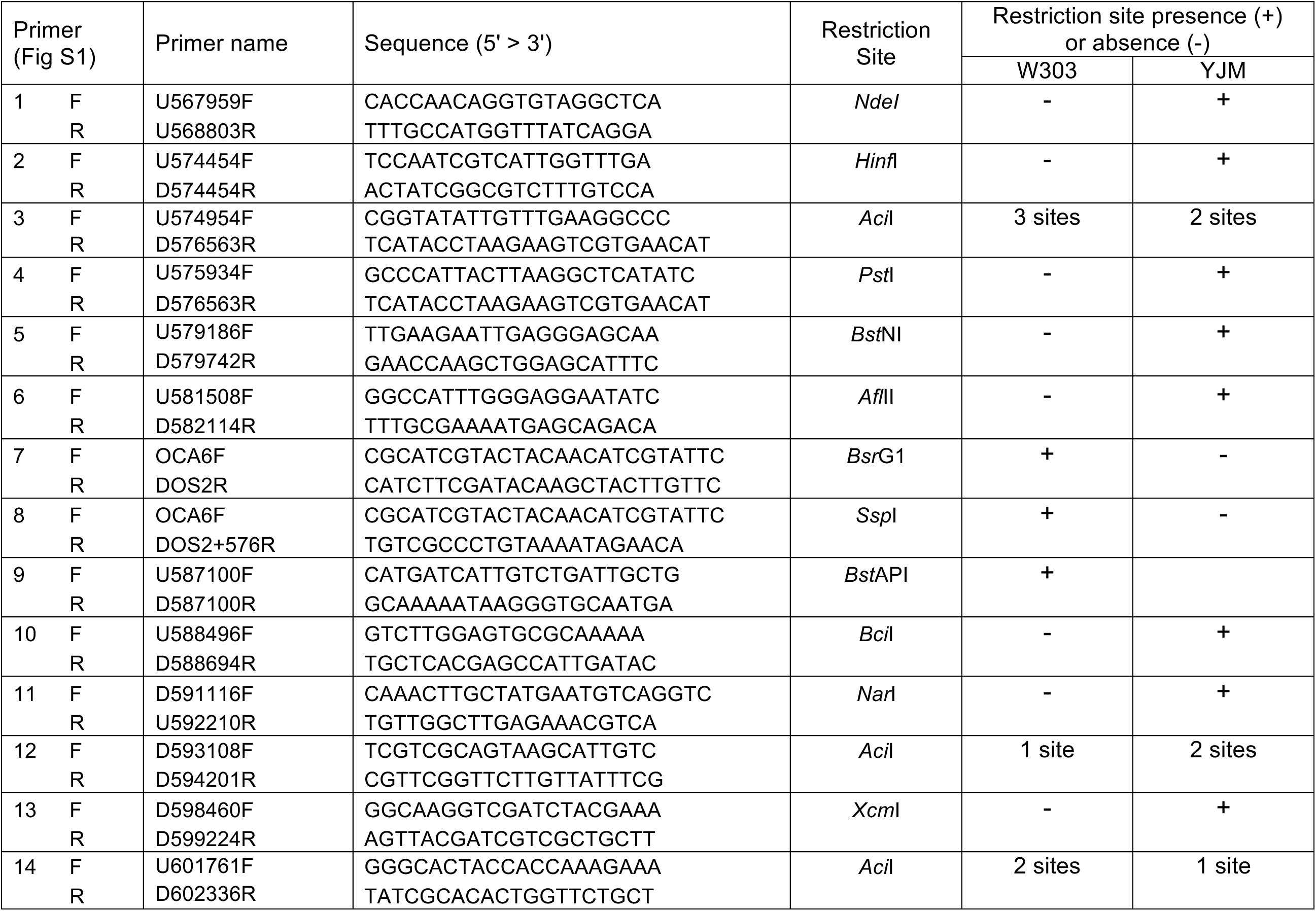
Primers and restriction site polymorphisms used to map GC tracts.

**Table S3.** Linkage relationships on CO and NCO chromosomes in the same sector.

| Sector | Primer pairs | # P spores | # NP spores | Linkage |
| --- | --- | --- | --- | --- |
| 1R | 6 and 11 | 2 | 49 | NP |
| 2R | 6 and 11 | 2 | 40 | NP |
| 3R | 2 and 8 | 0 | 53 | NP |
| 3W | 6 and 8 | 22 | 0 | P |
| 4R | 2 and 9 | 0 | 35 | NP |
| 6R | 1 and 6 | 47 | 0 | P |
| 7R | 1 and 9 | 24 | 2 | P |
| 8R<sup>1</sup> | 2 and 9 | 49 | 3 | P |
|  | 2 and 12 | 49 | 3 | P |
| 9R | 3 and 8 | 32 | 0 | P |
| 10R<sup>2</sup> | 1 and 4 | 0 | 33 | NP |
|  | 1 and 6 | 33 | 0 | P |
|  | 1 and 10 | 30 | 3 | P |
| 11R | 2 and 9 | 23 | 4 | P |
| 12R | 2 and 8 | 36 | 0 | P |
| 13R | 2 and 5 | 16 | 0 | P |
|  | 2 and 9 | 16 | 0 | P |
| 14R | 6 and 9 | 45 | 1 | P |
| 14W | 6 and 11 | 22 | 2 | P |
| 15R | 2 and 6 | 24 | 0 | P |
|  | 2 and 9 | 24 | 0 | P |
| 16R | 2 and 10 | 1 | 23 | NP |
| 17R | 1 and 8 | 28 | 0 | P |
| 17W | 2 and 11 | 24 | 0 | P |
|  | 2 and 14 | 24 | 0 | P |
| 18R | 6 and 11 | 29 | 0 | P |
| 19W | 1 and 3 | 46 | 0 | P |
| 20R | 2 and 6 | 32 | 0 | P |
|  | 2 and 11 | 32 | 0 | P |

<sup>1</sup>Fragment produced with primer pair 12 was sequenced to confirm transition point
<sup>2</sup>Single SNP at coordinate 575520 is homozygous

**Figure S1.**
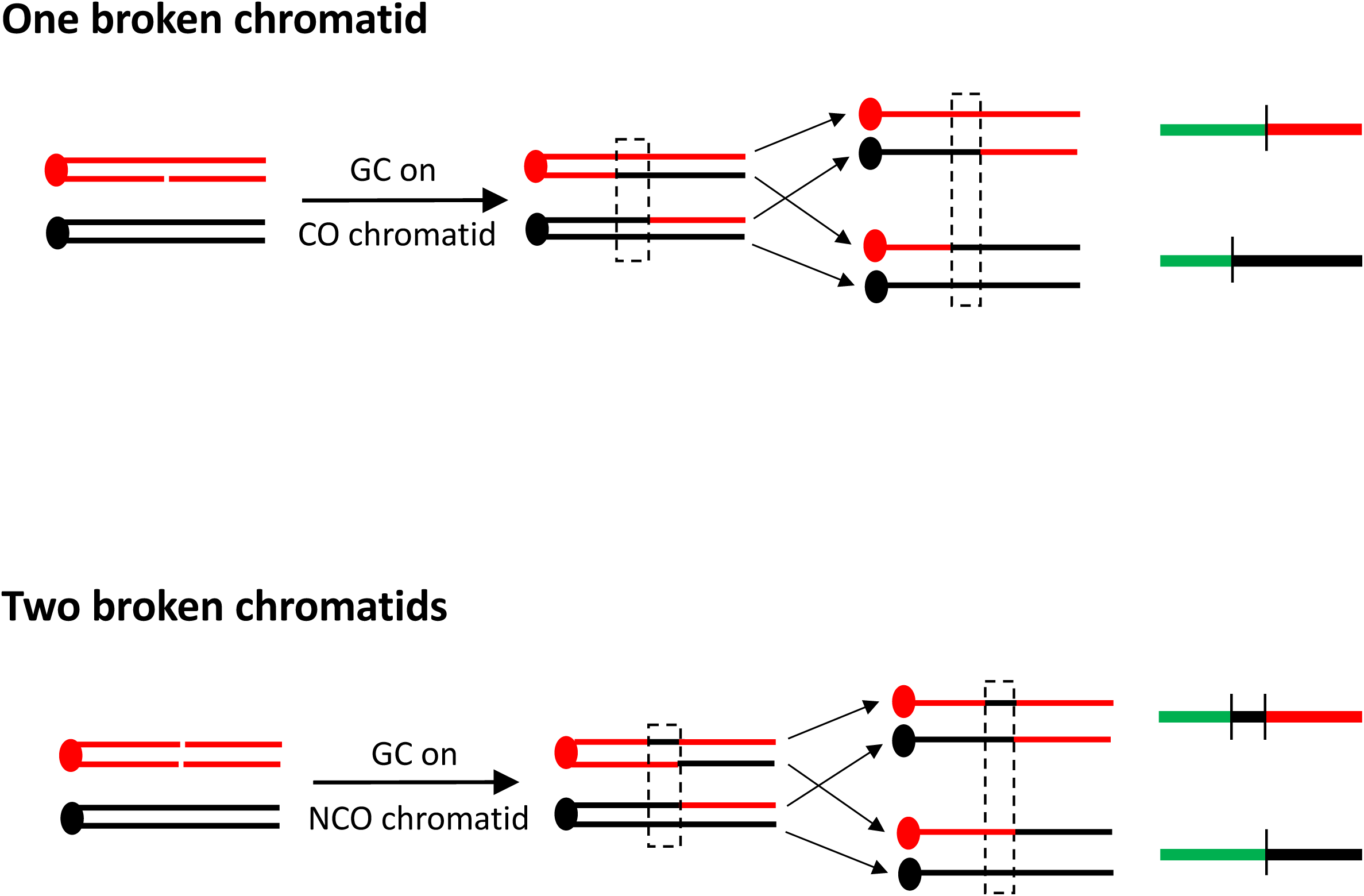
GC pattern when there are one versus two broken chromatids. Red and black lines correspond to W303-and YJM-specific chromosomal segments, respectively; red and black circles represent W303 and YJM centromeres, respectively. When only one chromatid is broken (top panel) the GC tract is always located on the CO chromatid with the W303 centromere. In the abbreviated red/black/green depiction (green indicates heterozygosity), there is a single transition between heterozygosity and homozygosity in each sector (green to red and green to black in the red and white sectors, respectively). When both chromatids are broken and repaired, however, the GC tract can be on the NCO, W303 chromatid instead of on the CO chromatid (lower panel). The segregation of the GC-containing, NCO chromatid with a CO chromatid leads to two transitions (green to black to red) in the red sector.

**Figure S2.**
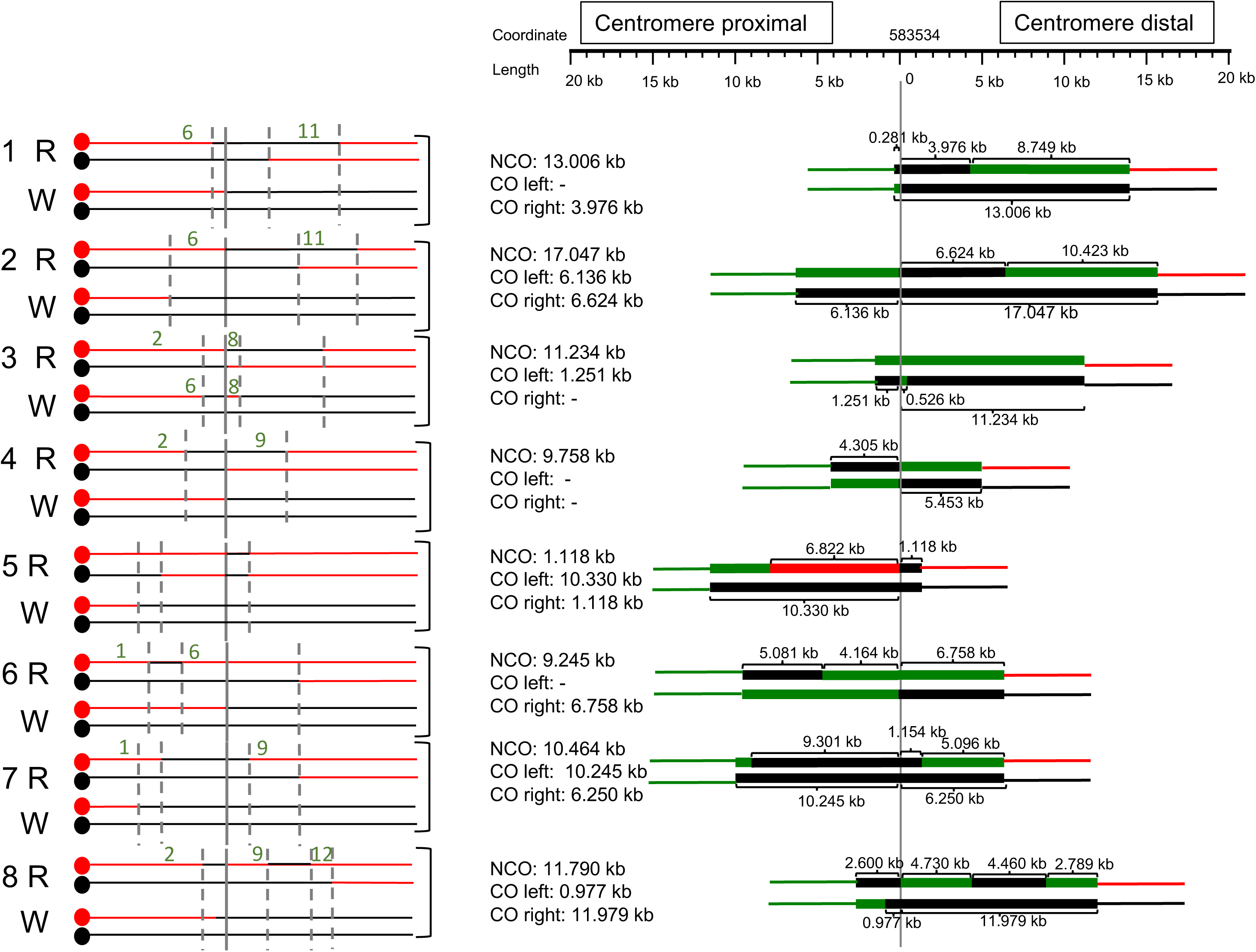
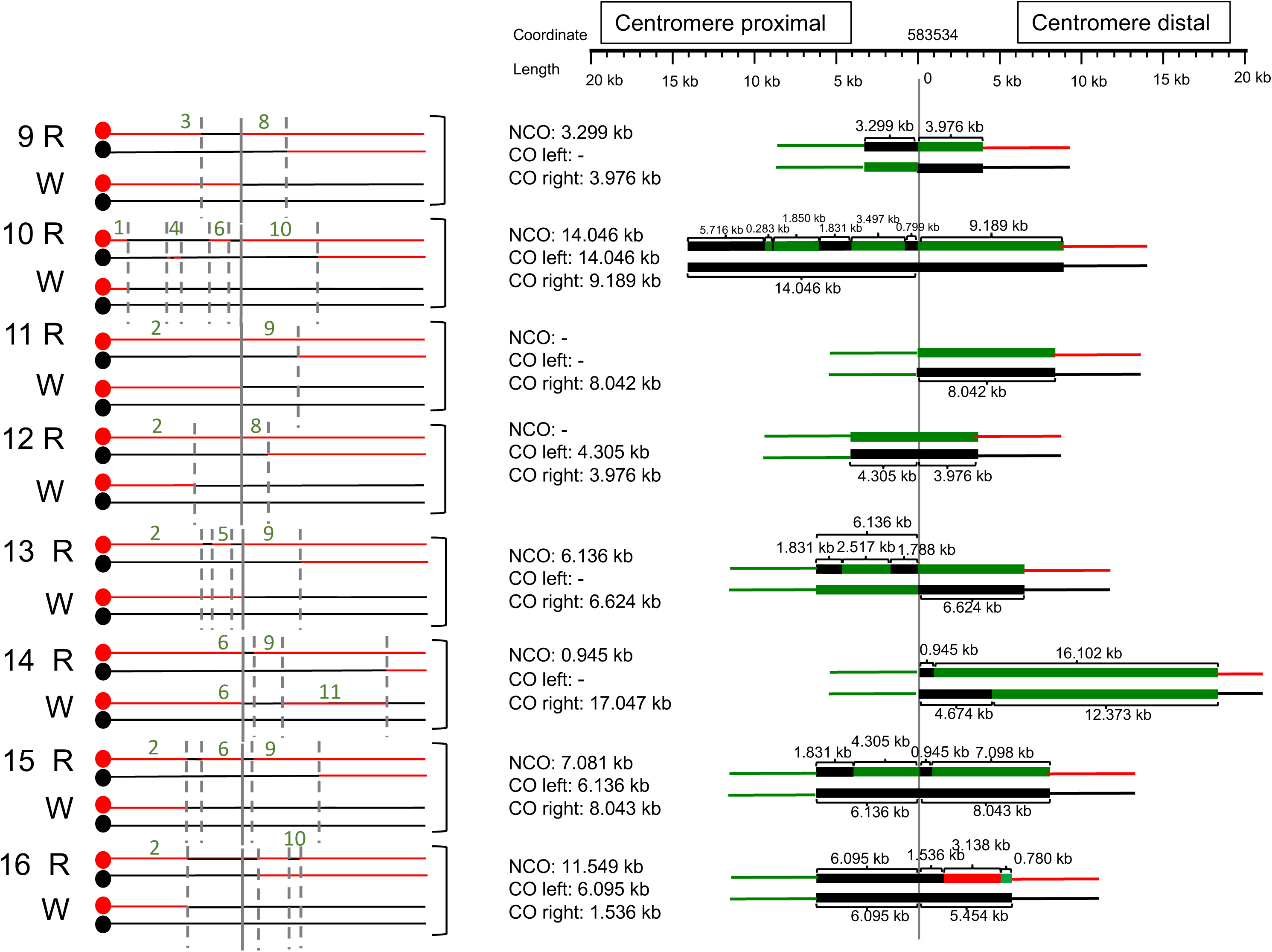
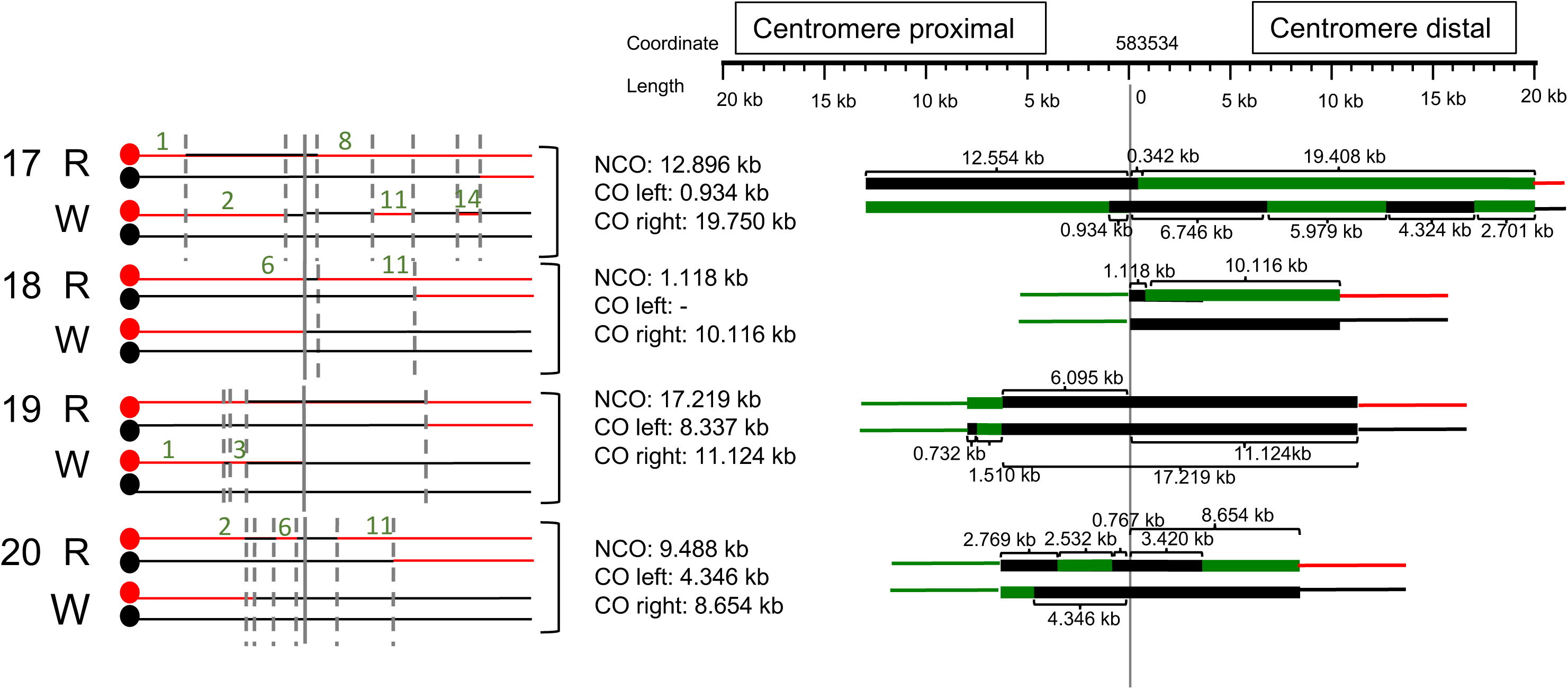
Chromosomes present in each sectored colony. The four chromosomes present within each sectored colony are diagrammed on the left. Red and black lines correspond to W303-and YJM-specific chromosomal segments, respectively, in CO and NCO products;; red and black circles represent W303 and YJM centromeres, respectively. Solid gray vertical lines mark the position of the initiating DSB and dotted lines mark transitions between heterozygosity and homozygosity. Green numbers correspond to primer pairs used to diagnose genetic linkages on the CO and NCO chromosomes within specific sectors (see Tables S2 and S3). On the right, the shorthand depiction is presented as in Figure 4, but 4:0 and 3:1 tract lengths are given and are presented to scale.

**Figure S3.**
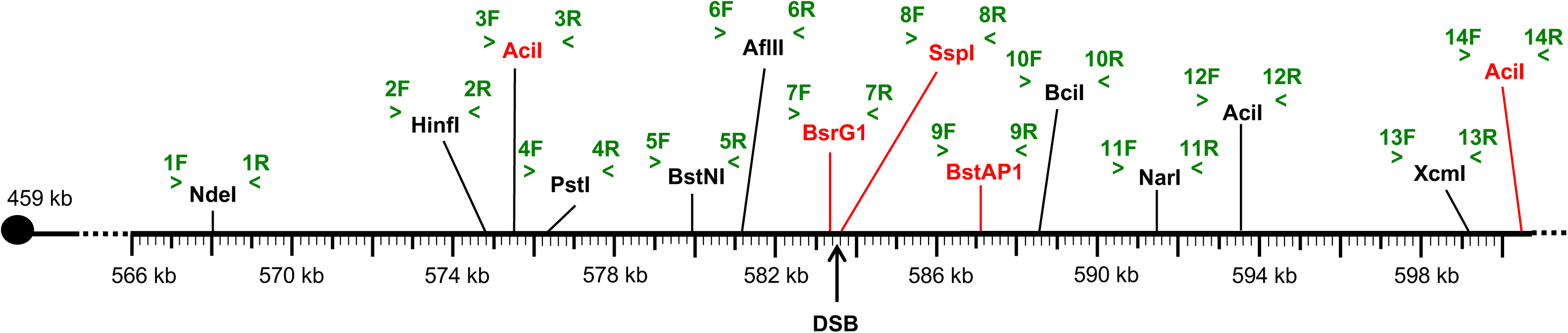
Primers pairs and restriction sites used to diagnose genetic linkages in spores. The right arm of chromosome IV is shown, with SGD coordinates indicated. Primer pairs are numbered and are in green. The W303-or YJM-specific restriction site within each corresponding PCR fragment is in red or black, respectively. The position of the initiating DSB is indicated.

